# The role of *Plasmodium falciparum* acyl-coenzyme A synthetase 10 in fatty acid acquisition and lipid synthesis

**DOI:** 10.64898/2026.11.06.764794

**Authors:** Katherine R. Fike, Halina Santos, Michael Klemba

**Affiliations:** Department of Biochemistry, Virginia Tech, Blacksburg, VA 24061

## Abstract

During the asexual blood stage, malaria parasites synthesize large quantities of lipids to support their rapid proliferation. Exogenous fatty acids obtained from host plasma are activated for lipid synthesis by parasite acyl-coenzyme A synthetases (ACSs). Of the twelve *Plasmodium falciparum* ACS homologs, ACS10 has emerged as a key blood-stage enzyme. Here, we sought to define the role of ACS10 in fatty acid acquisition and lipid synthesis during asexual development. To accomplish this, we have employed fatty acid alkyne probes of parasite lipid synthesis together with selective depletion of ACS10 in a conditional knockdown parasite line. This approach revealed that ACS10 is required for efficient incorporation of 16- to 20-carbon unsaturated fatty acid alkynes and a 20-carbon, saturated fatty acid alkyne into the major parasite phospholipids and neutral lipids. In contrast, shorter saturated fatty acid alkynes were either unaffected by ACS10 depletion or were hyper-incorporated into parasite lipids. ACS10 knockdown elicited a profound deformation of the food vacuole, which was likely caused by an imbalance of saturated and unsaturated fatty acids in parasite lipids. To promote the discovery and characterization of ACS inhibitors, we have established a fatty acid alkyne-based approach for identifying compounds that target ACS10 as their primary mode of action.

**IMPORTANCE:** Malaria is responsible for around 600,000 deaths annually, most which are caused by *Plasmodium falciparum*. The blood-stage parasite scavenges essential fatty acids from the host to support its vigorous lipid synthesis. ACSs serve as gateways of entry for fatty acids into the parasite’s lipid biosynthetic pathways and represent attractive therapeutic targets. We demonstrate that ACS10 plays a crucial role in recruiting essential fatty acids in support of parasite lipid synthesis. Loss of ACS10 activity had severe consequences for the parasite as exemplified by the dramatic distortion of a key organelle, the food vacuole. Having defined the metabolic contribution of ACS10, we developed a new approach for validating ACS10 inhibitors and thereby facilitating ACS-directed drug discovery.

## INTRODUCTION

Responsible for two million clinical cases and 600,000 deaths annually, malaria is a major public health burden that falls heavily on young children in sub-Saharan Africa (1). Artemisinin combination therapies are critical anti-malarial weapons for ameliorating this burden. While these remain generally effective in clinical practice, the appearance of parasite tolerance or resistance to artemisinin-family compounds and their partner drugs raises concerns over their long-term efficacy (2) and necessitates the development of anti-malarials with distinctive modes of action.

During the pathogenic, asexual erythrocytic stage, the malaria parasite relies on large quantities of host fatty acids to support the vigorous lipid synthesis needed for development of a single invasive merozoite into 16-32 progeny. Phosphatidylcholine (PC) and phosphatidylethanolamine (PE) constitute the major phospholipids at ∼55 and 35% of total phospholipids, respectively (3–5). Asexual parasites also produce substantial quantities of the neutral lipid triacylglycerol (TAG), which is deposited in lipid droplets that release free fatty acids towards the end of the replication cycle (6, 7).

Fatty acids are activated for lipid synthesis by thioesterification with coenzyme A, an ATP-dependent reaction that is catalyzed by the acyl-coenzyme A synthetase (ACS) family of enzymes. Two key themes have emerged from extensive studies of mammalian ACS isozymes. First, specific isozymes have distinctive specificities with respect to fatty acid length and degree of unsaturation (8). Second, the sub-cellular distributions of ACS isozymes can determine the metabolic fate of their acyl-CoA products, a phenomenon referred to as “fatty acid channeling” (9).

In the asexual malaria parasite, acyl-CoAs are essential substrates for *de novo* phospholipid and neutral lipid synthesis, phospholipid remodeling, and protein acylation but are not used as a source of metabolic energy. The genome of the most virulent human malaria parasite, *Plasmodium falciparum*, encodes 12 putatively functional ACS isozymes; four of these (ACS9-12) are conserved across the genus, and eight (ACS1a, 2-8; ACS1b is a pseudogene) represent a sub-telomeric expansion found in the great ape-infecting Laveranian clade (10, 11). Of the four genus-wide enzymes, ACS10 and 11 have attracted attention as candidate drug targets. Mutations in both have arisen in large-scale resistance-selection studies of anti-malarial compounds (12–15). ACS10 has been validated as a target of several anti-malarial chemotypes, while mutations in ACS11 appear to confer resistance by destabilizing the enzyme (15). Genetic knockdown has revealed that ACS10 is essential whereas ACS11 may not be (15). Lipidomic profiling of parasites in which ACS10 activity was depleted by knockdown or chemical inhibition have pointed to a role in lipid incorporation of 18-carbon unsaturated fatty acids and in promoting the synthesis of triacylglycerol (15). The development of drugs targeting ACSs is underway: the ACS inhibitor GSK701 (MMV1582367) is in the Medicines for Malaria Venture development pipeline and has recently completed phase I testing with favorable safety and pharmacokinetic profiles (16).

Given the relevance of ACS10 to anti-malarial discovery, a thorough understanding of its contributions to parasite lipid metabolism is desirable. On the basis of the lipidomic evidence in Bopp *et al* (15) and mentioned above, one possible hypothesis is that ACS10 channels a relatively limited subset of fatty acids (18 carbon, unsaturated) into a specific class of lipids (the neutral lipid triacylglycerol). We wished to further explore this hypothesis by investigating whether ACS10 contributes to: 1) the synthesis of the major parasite phospholipids PC and PE, thus indicating a broader role in supporting parasite lipid synthesis; and 2) incorporation of fatty acids other than unsaturated, 18-carbon fatty acids, to more completely define the structural characteristics of potential substrates. To accomplish these goals, we employed ω-alkyne analogs of fatty acids as probes of parasite fatty acid acquisition and lipid synthesis. Key features of fatty acid alkynes (FAA) that support their utility include minimal deviation from the structure of the natural fatty acid, high sensitivity after post-labeling derivatization, and convenience of use (17). We have previously shown that the ω-alkyne analog of oleic acid can serve as a reporter of lipid synthesis when added to the medium of cultured infected erythrocytes (18). Here, the role of ACS10 in promoting parasite lipid synthesis during asexual development was assessed by comparing the lipid incorporation of a panel of structurally-diverse FAAs in ACS10-replete and - depleted cultures. We took advantage of the insights thus gained to develop an approach for target validation of ACS10-directed inhibitors based on FAA profiling.

## RESULTS

### ACS10 is an essential enzyme located within the parasite

To enable the conditional depletion of ACS10, we installed a TetR-DOZI regulatory element at the 3’ end of the endogenous gene, either with or without the addition of a C-terminal hemagglutinin (HA) tag, through CRISPR gene editing (**Fig. S1**). Transfected parasites were cultured in the presence of anhydrotetracycline (aTc), which was permissive for ACS10 expression. Parasites were only obtained with constructs that maintained the native ACS C-terminus, which is consistent with a prior report (15) and implies that the unobstructed C-terminus plays an important structural or functional role. Withdrawal of anhydrotetracycline (-aTc; knockdown conditions) during early ring stage (0-4 h post-invasion) resulted in the disappearance of parasites on the following cycle (**Fig. 1A**), consistent with a previous report using the same knockdown system (15).

**Figure 1:**
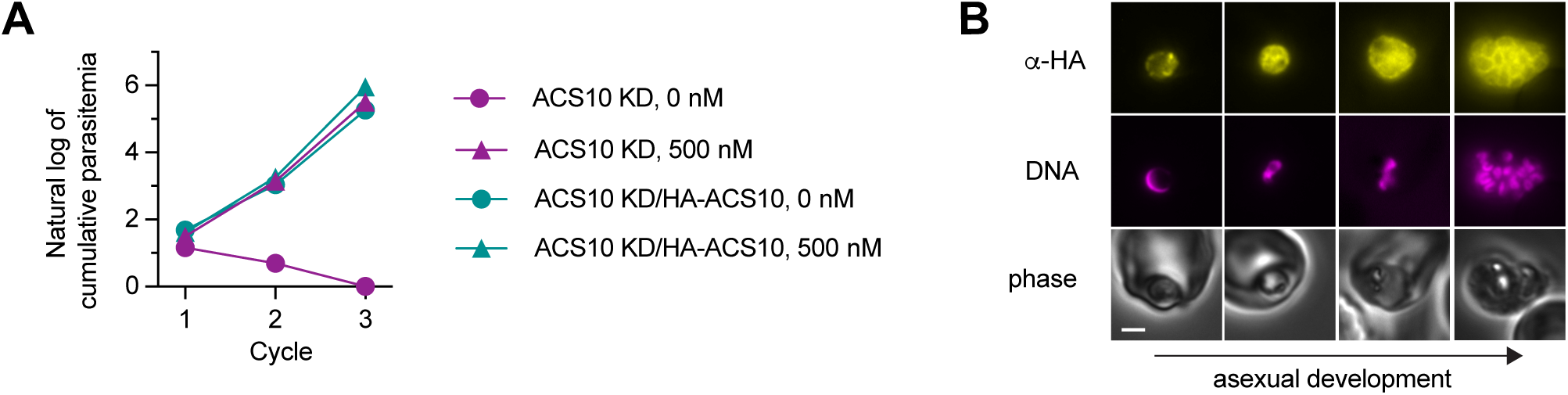
ACS10 is an essential enzyme that resides within the parasite. (A) Growth kinetics of ACS10 KD parasites under permissive (500 nM aTc) and knockdown (0 nM aTc) conditions, without and with complementation with HA-ACS10. Data for three 48-hour replication cycles are shown and are from a single experiment. (B) Immunofluorescence localization of HA-ACS10 across asexual development. Anti-HA fluorescence is pseudocolored yellow and Hoechst 33342 (DNA) fluorescence is pseudocolored magenta. Phase, brightfield phase contrast image. Scale bar, 2 µm.

To confirm that the lethality of withdrawing aTc was due to loss of ACS10, we complemented the knockdown line (termed ACS10 KD) with an N-terminal, triple hemagglutinin (3xHA)-tagged copy of ACS10 (HA-ACS10) expressed from a *piggybac* transposon, with transcription driven by the ACS10 promoter (**Fig. S1C**). Immunoblotting confirmed expression of HA-ACS10 (**Fig. S1D**). Growth of the ACS10 KD line expressing HA-ACS10 (termed ACS10 KD/HA-ACS10) was not affected by the removal of aTc (**Fig. 1A**), which unambiguously establishes the essential role of ACS10 and indicates that addition of an N-terminal epitope tag does not interfere with this role.

To define the cellular distribution of ACS10, HA-ACS10 was localized in asexual blood-stage parasites by immunofluorescence microscopy (**Fig. 1B**). The enzyme was observed within the parasite throughout the asexual cycle, with no evidence for export to the erythrocyte, which is consistent with the absence of a putative signal peptide at the N-terminus. Labeling of the parasite periphery was evident at all stages, suggesting association of HA-ACS10 with the parasite plasma membrane. Thus, ACS10 is situated to catalyze acyl-CoA formation at or near the parasite plasma membrane, thereby promoting a favorable concentration gradient for continuous inward diffusion of fatty acids, a phenomenon termed “vectorial acylation” (19). In trophozoites, labeling of internal features was also observed, which suggests an association with at least some organellar membranes.

### Fatty acid alkyne profiling of parasite lipid synthesis

To determine whether fatty acid alkynes (FAA) could be used in combination with ACS10 knockdown to characterize its role in providing acyl-CoAs for lipid synthesis, we assembled a panel of seven commercially-available FAA. Together, this panel provides diversity in both length and degree of unsaturation, representing ∼75% of the fatty acid composition of *P. falciparum* (**Table 1**). Fatty acid alkyne probes will be referred to throughout by reference to the natural fatty acid to which they correspond, *e.g.,* palmitate alkyne or 16:0 alkyne. Incorporation of 16:0 alkyne into the major parasite phospholipids, phosphatidylcholine (PC) and phosphatidylethanolamine (PE), was verified using authentic standards (**Fig. S2A**). No standard was available for triacylglycerol (TAG), but previous studies with fluorescent fatty acids have identified TAG as the highest-mobility (least polar) major labeled species in our TLC system (20). All of the probes in **Table 1** were efficiently incorporated into parasite lipids except for 16:1 alkyne, which was relatively weakly incorporated albeit at a level sufficient for quantitation (**Fig. S2B**).

**Table 1:**
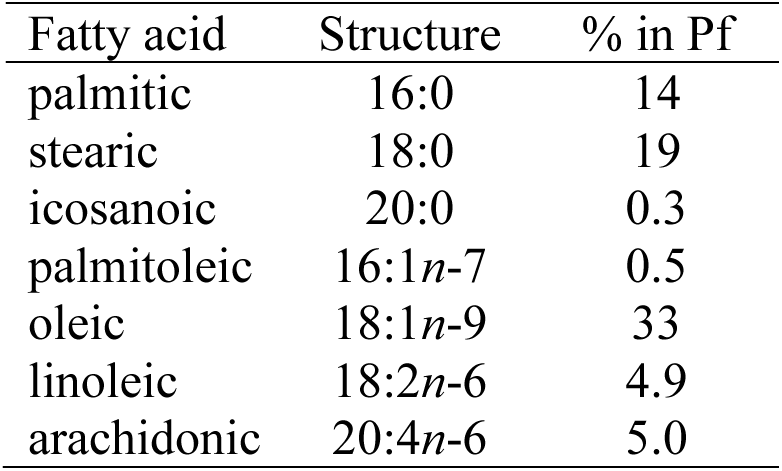
The relative abundance of fatty acids corresponding to ω-alkyne analogs used in this study in the asexual blood stage of *P. falciparum* (Pf). Values for % of total asexual *P. falciparum* fatty acids are from (15).

| Fatty acid | Structure | % in Pf |
|---|---|---|
| palmitic | 16:0 | 14 |
| stearic | 18:0 | 19 |
| icosanoic | 20:0 | 0.3 |
| palmitoleic | 16:1*n*-7 | 0.5 |
| oleic | 18:1*n*-9 | 33 |
| linoleic | 18:2*n*-6 | 4.9 |
| arachidonic | 20:4*n*-6 | 5.0 |

One saturated (18:0) and one unsaturated (18:1) FAA were selected for competition experiments to assess whether their uptake and incorporation into parasite lipids faithfully represents that of the corresponding natural fatty acid (FA). In each case, the FAA and FA were mixed in varying proportions with the total (FAA + FA) concentration maintained at 10 µM and were delivered in RPMI medium with fatty-acid free bovine serum albumin (BSA) as carrier along with an unsaturated fatty acid (for 18:0) or a saturated fatty acid (for 18:1) at 10 µM in order to provide the minimal saturated/unsaturated fatty acid mix that has been shown to support parasite growth (21). Trophozoite-stage ACS10 KD parasites (+500 nM aTc) were cultured in these media for one hour and incorporation of the alkyne into PC and PE was quantified. For 18:0 alkyne, a plot of relative alkyne labeling *vs*. fraction alkyne revealed slopes close to 1 (**Fig. 2A**, **Table 2**). Thus, 18:0 alkyne appears to be neither preferred nor disfavored relative to stearate. Slopes for 18:1 alkyne/oleate were somewhat lower at ∼ 0.7, which suggests that 18:1 alkyne is modestly disfavored relative to oleate. This may be a result of differences in binding affinities for BSA and/or any of the enzymes required to generate the final phospholipid products, or to differences in membrane diffusion rates. These results indicate that fatty acid alkynes are best utilized for relative quantitation, which in this study consisted of ACS10 KD parasites cultured under permissive and knockdown conditions.

**Figure 2:**
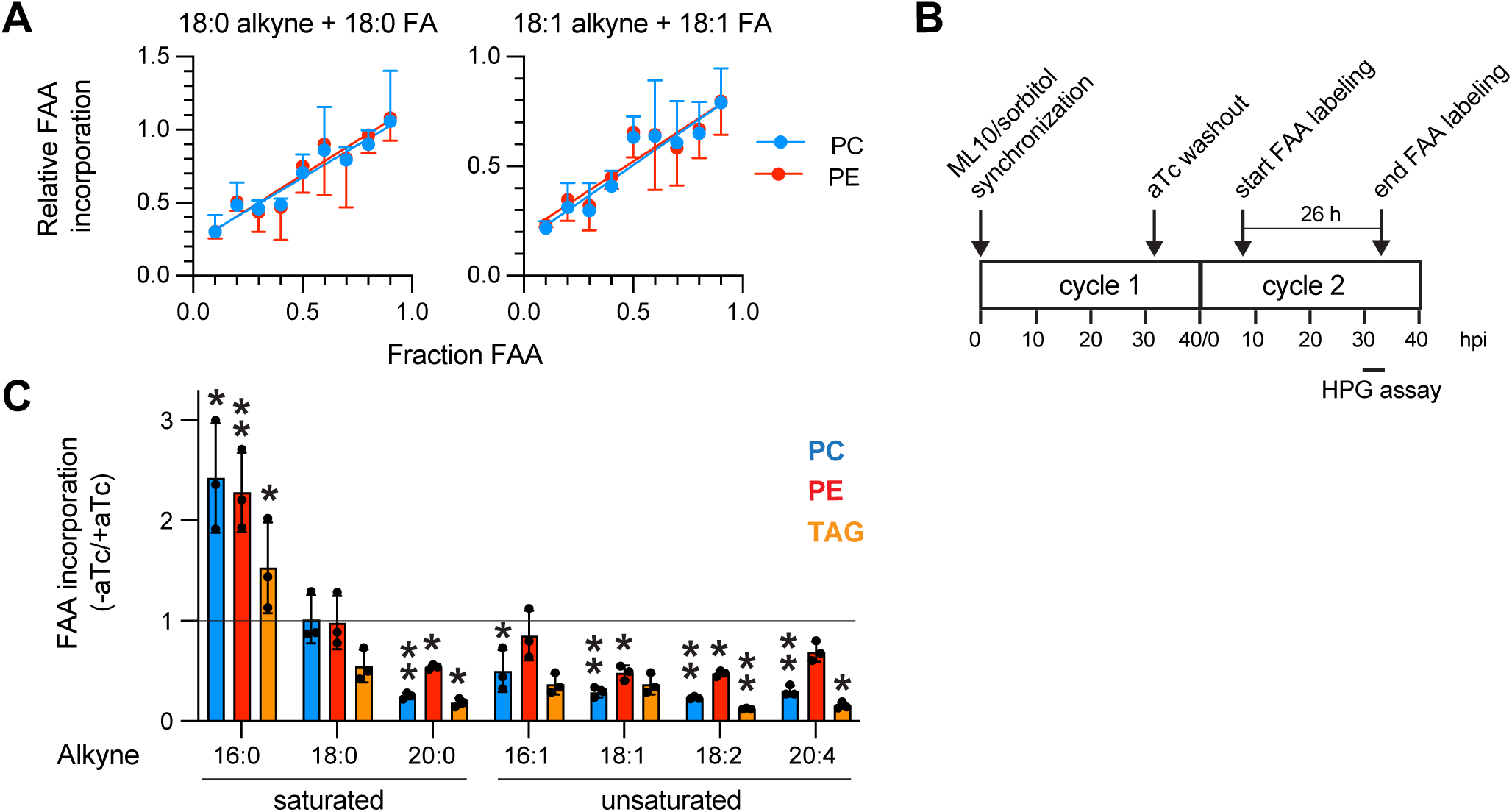
Fatty acid alkyne profiling of ACS10 knockdown parasites. (A) Competition experiments in which an FAA (18:0 or 18:1) was mixed with the corresponding fatty acid (FA) in 0.1 fractional increments. Data points are means ± SD of FAA incorporation into PC and PE, relative to the value for the alkyne-only control, from three independent experiments and are fitted by linear regression. (B) Schematic of the two-cycle workflow for ACS10 KD parasite synchronization, aTc washout, and FAA profiling. The timeframe over which protein synthesis assays (“HPG assay”) were conducted is indicated. (C) Effects of ACS10 knockdown on incorporation of seven FAA into PC, PE and TAG. Bars indicate mean FAA incorporation in knockdown (-aTc) parasites relative to that in permissive (+aTc) parasites from three independent experiments. The horizontal line indicates a value of 1, i.e., no effect of aTc knockdown. For each lipid, the significance of differences from the value for 18:0 alkyne were assessed with unpaired, two-tailed *t*-tests. One asterisk, p≤0.05; Two asterisks, p≤0.01.

**Table 2:**
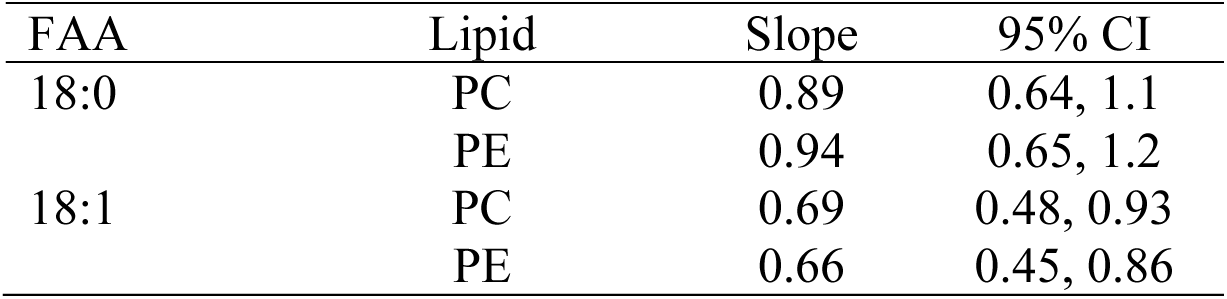
Slopes and 95% confidence intervals for FAA:FA competition experiments.

We sought to employ the FAA panel to profile the effects of ACS10 knockdown on lipid incorporation. For this to be effective, it was necessary to optimize the timing of aTc washout such that levels of ACS10 were as low as possible to maximize the effect of knockdown without causing metabolic collapse or parasite death. To identify an optimal workflow, parasites were synchronized to a <4 hour window with the egress inhibitor ML10 and sorbitol treatment. During the subsequent ∼40 hour replication cycle, the effects of aTc washout were assessed. We found that washout at 32 h post-invasion (hpi) permitted parasites to develop through the trophozoite stage during the following cycle; however, they failed to complete schizogony and re-invade efficiently. FAA labeling was therefore restricted to a 26-hour window during the post-washout cycle that terminated in the late trophozoite stage (see schematic in **Fig. 2B**). We were not able to quantify the extent of ACS10 knockdown achieved by this protocol due to the inability to introduce an epitope tag at the conditionally-regulated allele.

Each probe was supplied individually at a concentration of 10 µM in complete RPMI containing 0.5% Albumax I, which is a BSA-enriched supplement that provides exogenous fatty acids in the form of free fatty acids and lysophosphatidylcholine (LPC) (22). Thus, each FAA probe competed with the mixture of free fatty acids and LPC that exists under normal culturing conditions. Following the labeling period, alkyne-labeled lipids were extracted, conjugated to 3-azido-7-hydroxycoumarin by click chemistry, resolved by TLC, and imaged. Fluorescence associated with PC, PE and TAG was quantified, normalized to the internal standard BODIPY-TR-ceramide, and the effect of ACS10 knockdown was taken as the ratio of normalized fluorescence intensities in the presence and absence of aTc. To determine whether extended exposure to any of the fatty acid alkynes was detrimental to parasite viability, we sub-cultured an aliquot of each FAA-containing +aTc culture at the end of the labeling period and assessed re-growth after three days. There was no evidence for impaired proliferation in the presence of any FAA (**Fig. S2C**). To confirm the metabolic health of the cultures without aTc at the endpoint of the FAA labeling period (*i.e.*, 30-32 hpi), we quantified protein synthesis using the clickable methionine analog homopropargylglycine (HPG). Incorporation of HPG in -aTc cultures was 92 ± 14% of that in +aTc cultures (mean ± SD; n = 4 independent experiments), which indicates little, if any, loss of viability in the absence of aTc at this timepoint. The optimized workflow for FAA profiling and HPG analysis of ACS10 KD parasites is depicted in **Fig. 2B**.

The effects of ACS10 knockdown on FAA incorporation into parasite lipids varied with both probe length and presence of double bonds (**Fig. 2C, Fig. S2B**). 18:0 alkyne was used for statistical comparisons, as its incorporation into PC and PE was unaffected by ACS10 knockdown (mean -aTc:+aTc ratios of 1.0 and 0.98, respectively). In contrast to the case with phospholipids, incorporation of 18:0 alkyne into TAG was substantially reduced (mean -aTc:+aTc ratio of 0.55). This difference from the two phospholipids may be explained by the fact that TAG synthesis is initiated relatively late in the asexual cycle (7); thus, label incorporation into TAG may be more sensitive to small differences in development of the +aTc and -aTc cultures. Alternately, it may reflect a specific role for ACS10 in providing fatty acyl-CoA for the diacylglycerol acyltransferase-catalyzed synthesis of TAG from diacylglycerol. To maintain consistency with our analysis of PC and PE ratios, we used the 18:0 alkyne TAG ratio as the reference value for statistical comparisons.

Knockdown of ACS10 suppressed the incorporation of all unsaturated FAA probes into at least one of the three major parasite lipids (**Fig. 2C**). For all four probes, PC was significantly reduced, while PE and TAG were significantly reduced with two probes. No clear trends could be discerned with respect to unsaturated FAA probe length or degree of unsaturation. In contrast, the effect of ACS10 knockdown on lipid incorporation of saturated FAA probes was strongly dependent on length. Elongation of 18:0 alkyne by two carbons to 20:0 alkyne resulted in a sharp and significant reduction of incorporation into PC, PE and TAG to levels comparable to those of 20:4 alkyne. For these five probes, the magnitude of the effect of ACS10 knockdown was greater for PC than for PE (this difference was statistically significant for four probes; **Table 3**).

**Table 3:** Test of significance of differences in -aTc:+aTc FAA incorporation ratios for PC and PE. *p*-values are from two-tailed, unpaired *t*-tests.

| FAA | *p*-value |
|---|---|
| *Not significant* |  |
| 16:0 | 0.73 |
| 18:0 | 0.88 |
| 16:1 | 0.13 |
| *Significant* |  |
| 20:0 | 2.2 x 10<sup>-4</sup> |
| 18:1 | 2.0 x 10<sup>-2</sup> |
| 18:2 | 2.1 x 10<sup>-4</sup> |
| 20:4 | 4.0 x 10<sup>-3</sup> |

Shortening 18:0 alkyne by two carbons to 16:0 alkyne resulted in the opposite effect: ACS10 knockdown elicited a >2-fold increase in the 16:0 alkyne incorporation ratio for all three lipids relative to 18:0 alkyne.

### ACS10 knockdown distorts food vacuole and hemozoin morphology

In the course of these studies, we observed a highly aberrant morphology for the food vacuole hemozoin crystal in trophozoite-stage, ACS10-depleted parasites. Rather than appearing as a compact crystal or group of crystals, as observed in +aTc parasites, the hemozoin crystal frequently exhibited a needle-like, highly elongated morphology (**Fig. 3A**). We quantified the effect of ACS10 knockdown on hemozoin morphology by measuring the aspect ratio of the hemozoin crystal in stage-matched, early schizont (two-to four-nuclei) parasites. Parasites developing under ACS10 knockdown conditions exhibited a substantially higher mean aspect ratio compared to those cultured in the presence of aTc, an effect that was statistically significant in two independent replicates (**Fig. 3B**).

**Figure 3:**
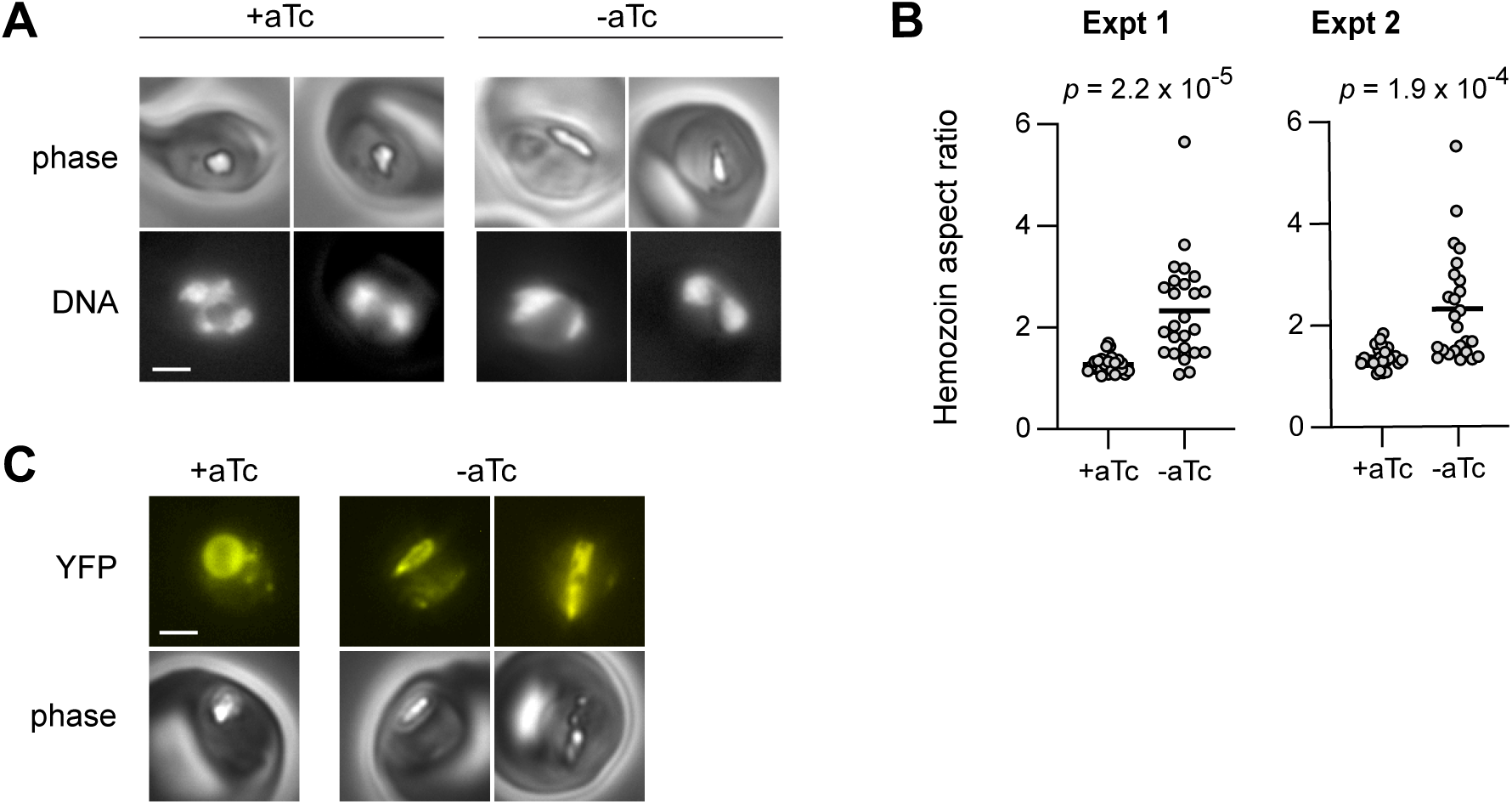
ACS10 knockdown induces aberrations in food vacuole and hemozoin morphology. (A) Images of stage-matched parasites cultured under permissive (+aTc) or knockdown (-aTc) conditions. The hemozoin crystal is the white feature within the parasite. (B) ACS10 knockdown increases the aspect ratio of the hemozoin crystal. Two independent experiments are shown, n=25 per condition. The mean value is indicated with a bar and the *p*-value from an unpaired, two-tailed Welch’s *t*-test is indicated. (C) Images of live ACS10 KD parasites expressing a plasmepsin II-YFP chimera, cultured with and without aTc. Scale bar, 2 µm.

To determine whether the food vacuole membrane was intact in parasites with aberrant hemozoin crystal morphology, we tagged the endogenous copy of the food vacuole aspartic protease plasmepsin II with yellow fluorescent protein (YFP) in the ACS10 KD line. Images of YFP fluorescence in live parasites cultured with aTc depicted a circular (in the two-dimensional image) food vacuole morphology (**Fig. 3C**), as expected (23). In parasites cultured without aTc and exhibiting elongated hemozoin crystals, YFP fluorescence was observed in close proximity to the crystal, indicating that the food vacuole membrane remained intact and was similarly distorted by ACS10 knockdown (**Fig. 3C**).

#### Quantifying in situ ACS10 inhibition by MMV019719

Given the potential clinical relevance of ACS-targeted drugs, we sought to adapt our profiling approach to enable identification of inhibitors targeting ACS10. For a test compound we selected MMV019719, which was identified as a putative ACS inhibitor by the appearance of a mutation in ACS11 upon selection for resistant parasites (12). Further investigation revealed that mutations at A268 of ACS10 could either sensitize (A268D) or confer resistance (A268V) to MMV019719 and that knockdown of ACS10 sensitized parasites (15). Interestingly, knockdown of ACS11 had the opposite effect of reducing sensitivity to MMV019719 (15). Together, these findings are consistent with ACS10 inhibition being the primary mode of action, with parasite resistance conferred by mutation or depletion of ACS11.

From the results of our FAA profiling experiments of ACS10 knockdown parasites, we reasoned that an ACS10-selective inhibitor would suppress lipid incorporation of 18:1 alkyne but not 16:0 alkyne. To avoid the confounding effects of long incubation periods with MMV019719 on parasite development and viability, we used a one-hour labeling assay with a simplified FAA/fatty acid composition. 18:1 alkyne was paired with palmitate (16:0), and 16:0 alkyne was paired with oleate (18:1), to provide the minimal saturated/unsaturated fatty acid mix that has been shown to support parasite growth (21). Cultures were incubated with a range of concentrations of MMV019719 for one hour with a subsequent one hour FAA labeling period. FAA incorporation into PC and PE was quantified. TAG was excluded from the analysis due to weak probe signal as a result of the shorter labeling period.

MMV019719 inhibited 18:1 alkyne incorporation into PC and PE in a concentration-dependent fashion, with IC_50_ values of 5.6 and 9.5 nM, respectively (**Fig. 4A**). The EC_50_ value for growth inhibition of the ACS10 KD parasite line (+aTc) was determined to be 150 ± 20 nM (mean ± SD, n = 4 independent experiments). Thus, it appears that a high level of inhibition of ACS10 is necessary to elicit an effect on parasite growth. MMV019719 did not induce a sigmoidal dose response curve for 16:0 alkyne, but rather a gradual, progressive decrease in incorporation into PC and PE (**Fig. 4B**). The absence of hyper-incorporation, as was observed in the FAA profiling experiments, is likely due to the much shorter (1 h *vs*. 26 h) time of exposure to probe in the MMV109719 assays. These findings provide direct support for a mode of action for MMV019719 based on inhibition of ACS10.

**Figure 4:**
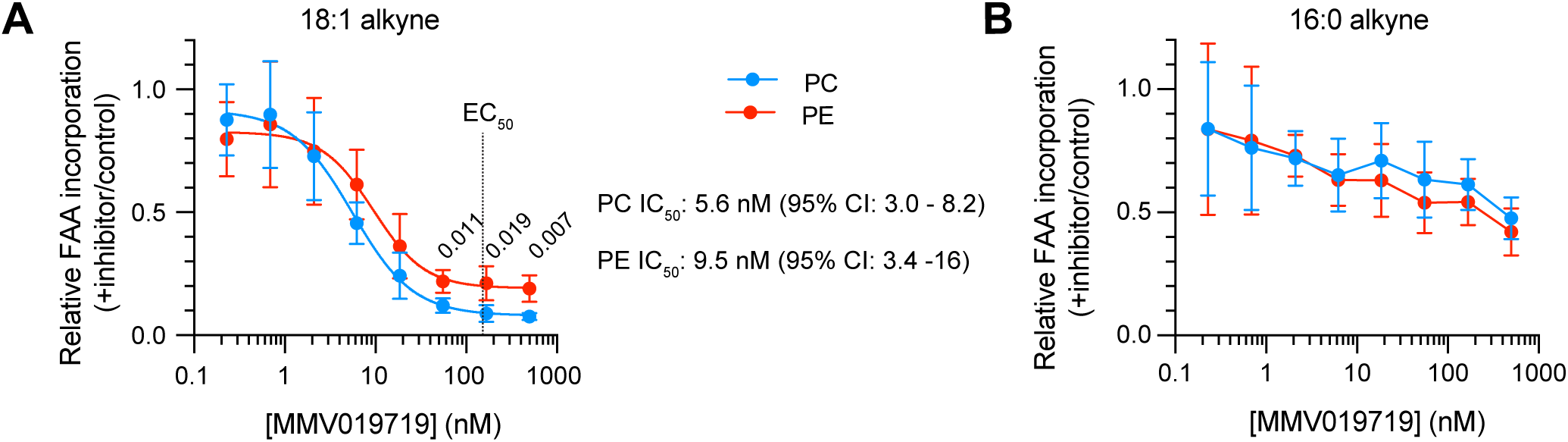
Effect of ACS10 inhibitor MMV019719 on incorporation of 18:1 alkyne (A) or 16:0 alkyne (B) into parasite PC and PE. Data points are means ± SD from four (A) and three (B) independent experiments. In (A), data points were fitted to a four-parameter concentration-response curve by non-linear regression. IC50 values and 95% confidence intervals for probe incorporation into each lipid are indicated. The EC50 value for growth inhibition by MMV019719 is indicated with a dashed vertical line. Numbers above selected data points are *p*-values from two-tailed *t*-tests. Data points in (B) were joined by lines to indicate a slight downward trend with increasing MMV019719 concentration.

Notably, the differential effect on PC and PE incorporation that was observed with ACS10 knockdown was recapitulated with the inhibitor: the fractional incorporation of 18:1 alkyne into PE remained approximately two-fold higher than that of PC at the three highest MMV019719 concentrations tested, at which point the lower baseline of the concentration-response curve is clearly established (**Fig. 4A**). This effect was observed for 18:1 alkyne but not 16:0 alkyne, which is consistent with expectation for ACS10 inhibition.

## DISCUSSION

We have described a new approach for interrogating the contributions of individual acyl-CoA synthetases to lipid synthesis that takes advantage of structurally-diverse FAAs and the ability to deplete a single ACS isozyme through conditional knockdown. This approach has the advantage of assessing the role of an ACS in its native physiological milieu in the intact organism and preserves the enzyme-membrane interactions that are likely to be an important factor in substrate selection and catalytic efficiency. While ACSs have been detergent-solubilized, purified, and characterized *in vitro*, the physiological relevance of the enzyme kinetic data emanating from such studies has been challenged on the basis of the disruption of enzyme-membrane interactions (24). FAA profiling offers an alternative approach and, as implemented here, provides a cumulative readout of the effects of ACS10 depletion on lipid incorporation across ∼24 hours of the ∼40 hour asexual replication cycle. While the observed effects of knockdown on FAA incorporation into lipids may be closely related to ACS10 substrate specificity, the latter cannot be conclusively inferred from the former due to the numerous enzyme-catalyzed steps separating fatty acyl (alkyne)-CoA formation from the final lipid products.

By expanding both the range of fatty acid structures and the classes of lipids impacted by ACS10 knockdown, our studies augment prior lipidomic analyses that identified 18-carbon unsaturated fatty acids as ACS10 putative substrates and triacylglycerol (TAG) as a lipid impacted by ACS10 depletion (15). Most significantly, we show that depletion of ACS10 reduces probe incorporation into PC to a similar extent to that of TAG. Interestingly, the magnitude of reduction of probe incorporation into PC was consistently greater than that into PE, both when ACS10 activity was depleted by knockdown and when it was inhibited by MMV019719.

Possible explanations include: 1) phospholipid remodeling alters the relative incorporation of unsaturated FAAs into PC and PE; or 2) synthesis of PE *via* decarboxylation of phosphatidylserine (25) in addition to the Kennedy pathway results in a different endpoint distribution of probe in this phospholipid. Further studies will be needed to understand the cause of this difference.

Resistance selection is a powerful tool for anti-malarial target identification; however, distinguishing mode of action from resistance mechanism can be challenging. Our FAA-based assay offers a direct and convenient approach to validation of ACS10-targeted compounds. The ACS10 inhibitor MMV019719 recapitulates the selectivity for 18:1 alkyne over 16:0 alkyne as well as the difference in PC and PE labeling that was observed with the ACS10 knockdown line. These studies suggest that relatively high fractional inhibition of probe incorporation (and possibly of ACS10 activity) is necessary to observe an effect on parasite growth.

## MATERIALS AND METHODS

### Key materials

16:1, 18:2 and 20:4 alkynes and 3-azido-7-hydroxycoumarin were from Cayman Chemical. 18:0, 18:1, 20:0 alkynes and 16:0(alkyne)-18:1-PC and 16:0(alkyne)-18:1-PE were from Avanti Polar Lipids. 16:0 alkyne, homopropargylglycine and CalFluor488 were from Vector Laboratories. BOIDPY-TR-ceramide (BTC) was from ThermoFisher.

### Parasite lines and culturing

*P. falciparum* clone 3D7 and ACS10 KD parasites were cultured in human O^+^ erythrocytes (Grifols Bio Supplies) at 2% hematocrit in RPMI 1640 medium supplemented with 0.37 mM hypoxanthine, 11 mM glucose, 27 mM sodium bicarbonate, 2 mM choline chloride, 10 μg/mL gentamicin and 5 g/L Albumax I (Gibco), unless otherwise stated. Cultures were incubated at 37 °C in a 5% CO_2_ with ambient O_2_ unless otherwise stated. For routine synchronization, the protein kinase G inhibitor ML10 (26) was used to arrest egress of segmented schizonts. Following ML10 washout, parasites were allowed to egress and invade for up to 4 hours, at which point the remaining schizonts were removed by treatment with 5% sorbitol. Methionine-free (Met-free) RPMI medium was made in-house. The construction of the new transgenic parasite lines is described in the Supplemental Material.

### Growth assays

Parasites were synchronized to a 4 h window using ML10 washout and sorbitol treatment and then split into +aTc and -aTc cultures. Parasitemia values over three cycles were quantified by counting >1000 erythrocytes on Giemsa stained smears.

### Immunoblotting and immunofluorescence

For immunoblotting, trophozoite-stage parasites (3D7 or ACS10KD/HA-ACS10) were harvested with saponin and lysed by sonication in PBS containing 5 µM pepstatin and 10 µM E-64 at a density of 6 x10^8^ parasites/mL. After rapid addition of SDS-PAGE sample buffer and heating to 95 °C for 5 minutes, samples were resolved on an 8.5% gel and blotted to nitrocellulose. The HA tag was detected using a rabbit anti-HA antibody (Invitrogen, #71-5500, 0.25 µg/mL) followed by an HRP-conjugated anti-rabbit IgG antibody.

For immunofluorescence, ACS10 KD/HA-ACS10 parasites cultured with 500 nM aTc were fixed in 2% paraformaldehyde and 0.0065% glutaraldehyde in PBS, permeabilized with 0.1% Triton-X100 and incubated with monoclonal anti-HA antibody 16B12 (Covance, 2 µg/mL) followed by an anti-mouse, Alexa488-conjugated antibody (2 µg/mL). DNA was visualized with Hoechst 33342 or DAPI. Images were collected on a Zeiss Axioimager M1 epifluorescence microscope with a 100x/1.4NA phase objective. Image contrast was adjusted with Adobe Photoshop.

### Fatty acid alkyne validation

Validation experiments were conducted with either: 1) 18:0 alkyne and 18:0 fatty acid at a total concentration of 10 µM, with the percentage of 18:0 alkyne ranging from 0 – 100% in 10% increments, and 10 µM 18:1 fatty acid; or 2) 18:1 alkyne and 18:1 fatty acid at a total concentration of 10 µM, in the same increments as in #1, and 10 µM 16:0 fatty acid. Test alkyne and fatty acid were each added separately at 10 µM to RPMI containing 3 mg/mL fatty acid-free bovine serum albumin and the complementing fatty acid. The resulting media were mixed to generate the desired FAA:fatty acid ratios.

A synchronized trophozoite culture of ACS10 KD clone B9 parasites was incubated with 1 µM BODIPY-TR-ceramide (BTC) for one hour at 37 °C in a CO_2_ incubator. Parasites were washed twice with incomplete RPMI, resuspended in the test media at 1 mL volumes, and incubated for one hour at 37 °C in a CO_2_ incubator. Cultures were washed once with complete RPMI and once with cold PBS. Lipids were extracted with methanol-chloroform and were subjected to click addition of 3-azido-7-hydroxycoumarin and TLC as previously described (18). BTC and probe fluorescence in PC and PE was imaged with an Azure 400 imager and quantified using ImageQuant software (Cytiva). Lipid fluorescence was normalized to that of the BTC internal standard.

### Fatty acid alkyne profiling

ACS10 KD clone B9 parasites cultured in the presence of 500 nM aTc were synchronized by blocking egress with ML10 followed by a washout period up to 4 hours to permit invasion. Remaining schizonts were removed by sorbitol treatment. At 32 hours post-invasion, parasites were washed four times with RPMI and split into -aTc and +aTc cultures. On the following day (∼48 hours post-invasion, or 8 hours into the second replication cycle), the cultures were collected by centrifugation and transferred to medium containing 10 µM of individual fatty acid alkynes in complete RPMI (containing 5 g/L Albumax I) and 0.2 µM BTC. After 24 hours (∼32 hours into the second replication cycle), a small aliquot of each +aTc culture was washed to remove the fatty acid alkyne and subcultured to assess viability by quantifying recovered parasitemia after three days. Labeled cultures were collected and fatty acid alkyne incorporation into PC, PE and TAG was quantified as described in “Fatty acid alkyne validation”. A schematic of this workflow is provided in **Fig. 2B**. A representative set of TLC images is provided in **Fig. S2C**.

### Protein synthesis assays

ACS10 KD clone B9 parasites cultured with and without 500 nM aTc were generated as described in “Fatty acid alkyne labeling” but without the addition of FAA to the medium. At 32 h into the second replication cycle (*i.e.*, when labeling reactions would be harvested), + and -aTc cultures were washed once in Met-free RPMI and taken up in Met-free RPMI containing 1 mM homopropargylglycine (HPG), without or with 5 µg/mL of the translation inhibitor cycloheximide to quantify background labeling. After 2 h incubation at 37 °C in a CO_2_ incubator, cultures were washed twice with complete RPMI and fixed with 0.1% glutaraldehyde. Proteins containing HPG were rendered fluorescent by click-addition of CalFluor488 and analyzed by flow cytometry as previously described (27). Protein synthesis was calculated by normalizing mean fluorescence intensities to those of solvent and cycloheximide controls. Means and standard deviations were calculated from four independent experiments.

### Hemozoin/food vacuole analysis

ACS10 KD clone B9 parasites cultured with and without 500 nM aTc were prepared as described in “Fatty acid alkyne labeling” but without the addition of FAA to the medium. At ∼32 h into the second replication cycle, parasites were fixed in 0.1% glutaraldehyde/PBS, adhered to polyethyleneimine-treated coverslips, and mounted in Prolong Diamond antifade mountant with DAPI (ThermoFisher). Parasites were stage-matched by limiting analysis to those with 2-4 nuclei as determined by DAPI staining. To ensure unbiased parasite selection, they were identified using DAPI fluorescence. Brightfield phase contrast images were obtained on a Zeiss AxioImager equipped with a 100x/1.4NA objective and a MRm Axiocam camera. Analysis of hemozoin crystal morphology was restricted to singly-infected erythrocytes in which a single, contiguous hemozoin crystal was visible. Aspect ratio was determined using Fiji/ImageJ by manually outlining a region around the hemozoin crystal. Twenty five parasites were analyzed per sample. Statistical significance was determined using an unpaired, two-tailed *t*-test.

Fluorescent images of live ACS10 KD parasites expressing a plasmepsin II-YFP fusion and cultured without or with 500 nM aTc were collected on a Zeiss Axioimager epifluorescence microscope equipped with a 100x/1.4NA objective.

### MMV019719 IC_50_ and EC_50_ assays

To determine IC_50_ values for incorporation of FAA probes into PC and PE, ACS10 KD clone B9 parasites were cultured with 500 nM aTc as described in “Fatty acid alkyne labeling” but without the addition of FAA to the medium. At ∼30 h into the second replication cycle, the culture was incubated with 1 µM BTC for one hour at 37 °C in a CO_2_ incubator. After two washes with incomplete RPMI, parasites were suspended in incomplete RPMI + 500 nM aTc at 4% hematocrit and aliquoted at 500 µL volumes into a 24 well plate. MMV019719 was added from 1000x DMSO stocks to generate a 2-fold dilution series (range: 500 – 0.23 nM). After one hour incubation at 37 °C in a CO_2_ incubator, 500 µL of RPMI + MMV019719 at 1x concentration + 6 mg/mL BSA + 20 µM each of fatty acid alkyne and fatty acid (*i.e.*, either 18:1 alkyne + 16:0 fatty acid, or 16:0 alkyne + 18:1 fatty acid) were transferred to the culture, yielding final concentrations of 3 mg/mL BSA and 10 µM of the fatty acid alkyne and paired fatty acid. After one hour at 37 °C in a CO_2_ incubator, labeled cultures were collected and fatty acid alkyne incorporation into PC and PE was analyzed as described in “Fatty acid alkyne validation”. Values were normalized to those obtained with vehicle-only (0.1% DMSO) controls. EC_50_ values for inhibition of parasite replication were conducted with synchronized ring-stage cultures incubated with a two-fold concentration series of MMV019719 (range: 4.8 µM to 4.7 nM) for 48 hours in an atmosphere of 5% O_2_, 5% CO_2_ and 90% N_2_. SYBR Green quantitation of parasite nucleic acids was conducted as previously described (28). To obtain IC_50_ and EC_50_ values, data points were fitted to a four-parameter sigmoidal concentration-response equation.

## Acknowledgments

We are grateful to Christiaan van Ooij and LifeArc for ML10, Benoît Laleu and Medicines for Malaria Venture for MMV019719, and Jacobus Pharmaceuticals for WR99210. This work was supported by National Institutes of Health grant R21AI78339 and Virginia Agricultural Experiment Station project VA-137938 (M.K.). The content is solely the responsibility of the authors and does not necessarily represent the official views of the National Institutes of Health.

## Data availability

All data are contained within the manuscript.

## REFERENCES

1. World malaria report 2025: addressing the threat of antimalarial drug resistance. World Health Organization.

2. Martin RE, Shafik SH, Richards SN. 2018. Mechanisms of resistance to the partner drugs of artemisinin in the malaria parasite. Curr Opin Pharmacol 42:71–80.

3. Tran PN, Brown SH, Rug M, Ridgway MC, Mitchell TW, Maier AG. 2016. Changes in lipid composition during sexual development of the malaria parasite *Plasmodium falciparum*. Malar J 15:73.

4. Gulati S, Ekland EH, Ruggles KV, Chan RB, Jayabalasingham B, Zhou B, Mantel PY, Lee MC, Spottiswoode N, Coburn-Flynn O, Hjelmqvist D, Worgall TS, Marti M, Di Paolo G, Fidock DA. 2015. Profiling the essential nature of lipid metabolism in asexual blood and gametocyte stages of *Plasmodium falciparum*. Cell Host Microbe 18:371–81.

5. Vial HJ, Ancelin ML, Thuet MJ, Philippot JR. 1989. Phospholipid metabolism in *Plasmodium*-infected erythrocytes: guidelines for further studies using radioactive precursor incorporation. Parasitology 98:351–7.

6. Jackson KE, Klonis N, Ferguson DJ, Adisa A, Dogovski C, Tilley L. 2004. Food vacuole-associated lipid bodies and heterogeneous lipid environments in the malaria parasite, *Plasmodium falciparum*. Mol Microbiol 54:109–22.

7. Palacpac NM, Hiramine Y, Mi-ichi F, Torii M, Kita K, Hiramatsu R, Horii T, Mitamura T. 2004. Developmental-stage-specific triacylglycerol biosynthesis, degradation and trafficking as lipid bodies in Plasmodium falciparum-infected erythrocytes. J Cell Sci 117:1469–80.

8. Soupene E, Kuypers FA. 2008. Mammalian long-chain acyl-CoA synthetases. Exp Biol Med (Maywood) 233:507–21.

9. Mashek DG, Li LO, Coleman RA. 2007. Long-chain acyl-CoA synthetases and fatty acid channeling. Future Lipidol 2:465–476.

10. Bethke LL, Zilversmit M, Nielsen K, Daily J, Volkman SK, Ndiaye D, Lozovsky ER, Hartl DL, Wirth DF. 2006. Duplication, gene conversion, and genetic diversity in the species-specific acyl-CoA synthetase gene family of *Plasmodium falciparum*. Mol Biochem Parasitol 150:10–24.

11. Matesanz F, Tellez M, Alcina A. 2003. The *Plasmodium falciparum* fatty acyl-CoA synthetase family (PfACS) and differential stage-specific expression in infected erythrocytes. Mol Biochem Parasitol 126:109–12.

12. Cowell AN, Istvan ES, Lukens AK, Gomez-Lorenzo MG, Vanaerschot M, Sakata-Kato T, Flannery EL, Magistrado P, Owen E, Abraham M, LaMonte G, Painter HJ, Williams RM, Franco V, Linares M, Arriaga I, Bopp S, Corey VC, Gnadig NF, Coburn-Flynn O, Reimer C, Gupta P, Murithi JM, Moura PA, Fuchs O, Sasaki E, Kim SW, Teng CH, Wang LT, Akidil A, Adjalley S, Willis PA, Siegel D, Tanaseichuk O, Zhong Y, Zhou Y, Llinas M, Ottilie S, Gamo FJ, Lee MCS, Goldberg DE, Fidock DA, Wirth DF, Winzeler EA. 2018. Mapping the malaria parasite druggable genome by using *in vitro* evolution and chemogenomics. Science 359:191–199.

13. Summers RL, Pasaje CFA, Pisco JP, Striepen J, Luth MR, Kumpornsin K, Carpenter EF, Munro JT, Lin, Plater A, Punekar AS, Shepherd AM, Shepherd SM, Vanaerschot M, Murithi JM, Rubiano K, Akidil A, Ottilie S, Mittal N, Dilmore AH, Won M, Mandt REK, McGowen K, Owen E, Walpole C, Llinas M, Lee MCS, Winzeler EA, Fidock DA, Gilbert IH, Wirth DF, Niles JC, Baragana B, Lukens AK. 2022. Chemogenomics identifies acetyl-coenzyme A synthetase as a target for malaria treatment and prevention. Cell Chem Biol 29:191–201.

14. Schalkwijk J, Allman EL, Jansen PAM, de Vries LE, Verhoef JMJ, Jackowski S, Botman PNM, Beuckens-Schortinghuis CA, Koolen KMJ, Bolscher JM, Vos MW, Miller K, Reeves SA, Pett H, Trevitt G, Wittlin S, Scheurer C, Sax S, Fischli C, Angulo-Barturen I, Jimenez-Diaz MB, Josling G, Kooij TWA, Bonnert R, Campo B, Blaauw RH, Rutjes F, Sauerwein RW, Llinas M, Hermkens PHH, Dechering KJ. 2019. Antimalarial pantothenamide metabolites target acetyl-coenzyme A biosynthesis in *Plasmodium falciparum*. Sci Transl Med 11:eaas9917.

15. Bopp S, Pasaje CFA, Summers RL, Magistrado-Coxen P, Schindler KA, Corpas-Lopez V, Yeo T, Mok S, Dey S, Smick S, Nasamu AS, Demas AR, Milne R, Wiedemar N, Corey V, Gomez-Lorenzo MG, Franco V, Early AM, Lukens AK, Milner D, Furtado J, Gamo FJ, Winzeler EA, Volkman SK, Duffey M, Laleu B, Fidock DA, Wyllie S, Niles JC, Wirth DF. 2023. Potent acyl-CoA synthetase 10 inhibitors kill *Plasmodium falciparum* by disrupting triglyceride formation. Nat Commun 14:1455.

16. Kuemmerle A, Singh N, Gossen D, Janin A, Sharma R, Cahn A, Gibson RA, Menakuru SR, Lambourne E, Dove T, Gamo FJ, Sanz L, Bestgen B, Chalon S. 2025. First-in-human safety, tolerability, pharmacokinetics and pilot food-effect study of the candidate antimalarial compound MMV367. Br J Clin Pharmacol 91:1821–1833.

17. Thiele C, Papan C, Hoelper D, Kusserow K, Gaebler A, Schoene M, Piotrowitz K, Lohmann D, Spandl J, Stevanovic A, Shevchenko A, Kuerschner L. 2012. Tracing fatty acid metabolism by click chemistry. ACS Chem Biol 7:2004–11.

18. Liu J, Fike KR, Dapper C, Klemba M. 2024. Metabolism of host lysophosphatidylcholine in *Plasmodium falciparum*-infected erythrocytes. Proc Natl Acad Sci USA 121:e2320262121.

19. Black PN, DiRusso CC. 2007. Vectorial acylation: linking fatty acid transport and activation to metabolic trafficking. Novartis Found Symp 286:127–38.

20. Dapper C, Liu J, Klemba M. 2022. Leveraging a fluorescent fatty acid probe to discover cell-permeable inhibitors of *Plasmodium falciparum* glycerolipid biosynthesis. Microbiol Spectr 10:e0245622.

21. Mi-Ichi F, Kano S, Mitamura T. 2007. Oleic acid is indispensable for intraerythrocytic proliferation of *Plasmodium falciparum*. Parasitology 134:1671–7.

22. Garcia-Gonzalo FR, Izpisua Belmonte JC. 2008. Albumin-associated lipids regulate human embryonic stem cell self-renewal. PLoS One 3:e1384.

23. Klemba M, Beatty W, Gluzman I, Goldberg DE. 2004. Trafficking of plasmepsin II to the food vacuole of the malaria parasite *Plasmodium falciparum*. J Cell Biol 164:47–56.

24. Soupene E, Fyrst H, Kuypers FA. 2008. Mammalian acyl-CoA:lysophosphatidylcholine acyltransferase enzymes. Proc Natl Acad Sci USA 105:88–93.

25. Baunaure F, Eldin P, Cathiard AM, Vial H. 2004. Characterization of a non-mitochondrial type I phosphatidylserine decarboxylase in *Plasmodium falciparum*. Mol Microbiol 51:33–46.

26. Ressurreicao M, Thomas JA, Nofal SD, Flueck C, Moon RW, Baker DA, van Ooij C. 2020. Use of a highly specific kinase inhibitor for rapid, simple and precise synchronization of *Plasmodium falciparum* and *Plasmodium knowlesi* asexual blood-stage parasites. PLoS One 15:e0235798.

27. Shieh P, Dien VT, Beahm BJ, Castellano JM, Wyss-Coray T, Bertozzi CR. 2015. CalFluors: A universal motif for fluorogenic azide probes across the visible spectrum. J Am Chem Soc 137:7145–51.

28. Smilkstein M, Sriwilaijaroen N, Kelly JX, Wilairat P, Riscoe M. 2004. Simple and inexpensive fluorescence-based technique for high-throughput antimalarial drug screening. Antimicrob Agents Chemother 48:1803–6.

